# Concurrent single-pulse (sp) TMS/fMRI to reveal the causal connectome in healthy and patient populations

**DOI:** 10.1101/2024.09.25.614833

**Authors:** Cameron Glick, Niharika Gajawelli, Yinming Sun, Faizan Badami, Manish Saggar, Amit Etkin

## Abstract

Neuroimaging and cognitive neuroscience studies have identified neural circuits linked to anxiety, mood, and trauma-related symptoms and focused on their interaction with the medial prefrontal default mode circuitry. Despite these advances, developing new neuromodulatory treatments based on neurocircuitry remains challenging. It remains unclear which nodes within and controlling these circuits are affected and how their impairment is connected to psychiatric symptoms. Concurrent single-pulse (sp) TMS/fMRI offers a promising approach to probing and mapping the integrity of these circuits. In this study, we present concurrent sp-TMS/fMRI data along with structural MRI scans from 152 participants, including both healthy and depressed individuals. The sp-TMS was administered to 11 different cortical sites, providing a dataset that allows researchers to investigate how brain circuits are modulated by spTMS.

## Background & Summary

The human brain at rest is inherently organized into large-scale functional networks (e.g., Power et al., 2011; Deco, Jirsa, and McIntosh, 2011), which are thought to play a crucial role in the transmission of information between brain regions (Cole et al., 2016). While there is substantial literature on the complex dynamics within and between these functional networks (e.g., Misic et al., 2015; Gollo, Roberts, and Cocchi, 2017), how these intrinsic networks influence or cause activity flow in the human brain remains unclear due to a lack of direct empirical evidence on the propagation of local activity across the brain. This gap in our understanding, essential for uncovering the causal neural mechanisms underlying brain functions, can potentially be addressed by integrating transcranial magnetic stimulation (TMS) and functional magnetic resonance imaging (fMRI). TMS has long been used to manipulate brain activity in cognitive neuroscience and therapeutic contexts. Each TMS pulse generates a brief magnetic field that penetrates biological tissue, inducing an electric field in the targeted brain area, typically affecting about 1 cm^3^ of the superficial cortex. Significantly, TMS not only influences neural activity at the stimulation site but also triggers changes in distal, interconnected brain regions (Bestmann, Ruff, Blankenburg, et al., 2008; Ruff, Driver, and Bestmann, 2009). This suggests that observed activity changes in non-stimulated regions could result from activity propagation through intrinsic connectivity from the stimulated site.

Single TMS pulses can probe the integrity of these circuits, while repetitive TMS (rTMS) can modulate cortical activity—high-frequency rTMS enhances activity, while low-frequency rTMS suppresses it. The primary goal of a single rTMS session is to determine if it can temporarily improve brain abnormalities in patients and be used as a target for future interventions rather than to create lasting therapeutic changes.

By integrating TMS with concurrent fMRI (TMS/fMRI), we can visualize how localized TMS affects widespread downstream neural circuits. This combined approach offers insights into how different prefrontal TMS nodes regulate these circuits and whether such effects differ in patients with anxiety, depression, or trauma-related symptoms—populations that have been unexplored in concurrent TMS/fMRI studies.

We provide a comprehensive TMS/fMRI dataset that includes resting-state recordings and single-pulse TMS-fMRI data. This dataset enables the exploration of cortical activity changes induced by spTMS at 11 different sites in both trauma-exposed and non-trauma-exposed healthy participants, as well as in trauma-induced and non-trauma symptomatic depressed participants. The resting-state data can be used to assess changes in functional connectivity metrics, while the spTMS data allows for investigating spatiotemporal brain changes across different stimulation sites.

## Methods

### Participant Population

Participants were asked to participate in 4 study visits, the first of which included consent, intake, and neuropsychological test; the second: anatomical T1, DWI, and fMRI data acquisition; the third: TMS/fMRI scans; the fourth: TMS/EEG. This dataset reports subset data collected from the second and third study visits (**Figure 2A**). Due to constraints on scan time, intolerability of specific stimulation sites for some participants, unusable noisy data, and the addition of some targets after the start of the study, not all participants completed all TMS/fMRI runs.

All participants provided signed consent, and their participation in the study was monitored by trained study staff. The study was approved by the Stanford University Institutional Review Board. Participants were excluded if they demonstrated or reported any significant neurological pathology, e.g., loss of consciousness >30 mins, post-trauma amnesia >24 hrs. Furthermore, exclusion criteria included claustrophobia and the use of psychiatric medication. Doctoral-level clinicians administered the *Structured Clinical Interview for DSM-IV Disorders* to characterize current and past psychiatric pathologies in participants. Participants were then grouped into 4 cohorts, each representing a subsample of the study population. The cohorts are as follows: Non-Trauma Healthy Control (NTHC), Trauma Exposed Healthy Control (TEHC), Non-Trauma Symptomatic (NTS), and Trauma-Induced Symptomatic (TIS) (**Figure 1A**). Throughout the study, participants were encouraged to report any physical or psychological discomfort and could withdraw at any time.

**Fig 1:**
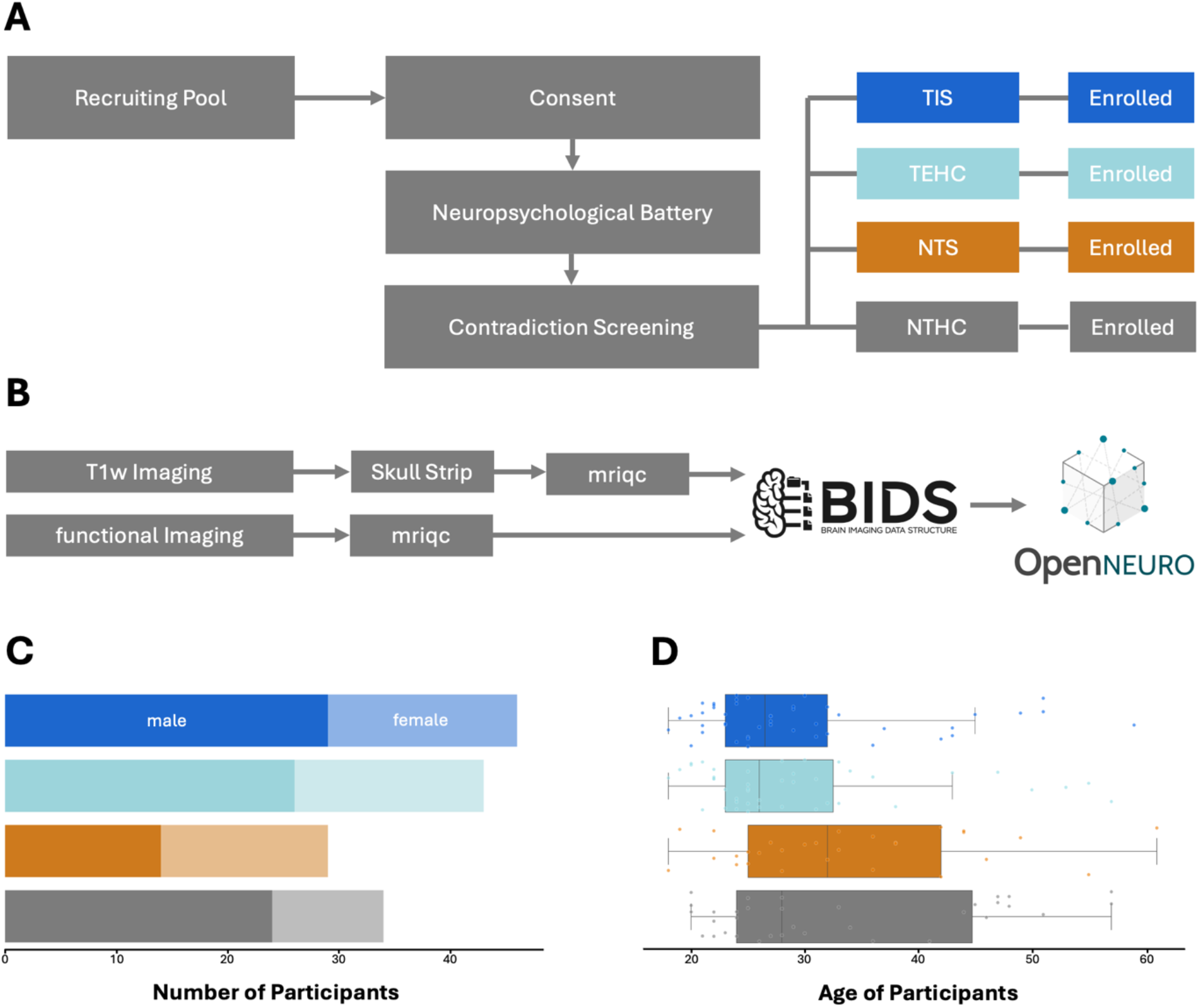
(A) Recruitment Pipeline. Every participant undergoes a screening process and is then separated into one of 4 groups. (B) Imaging pipeline. (C) Gender balance per group for all reported participants. (D) Age per group (in years) for all reported participants.

**Fig 2:**
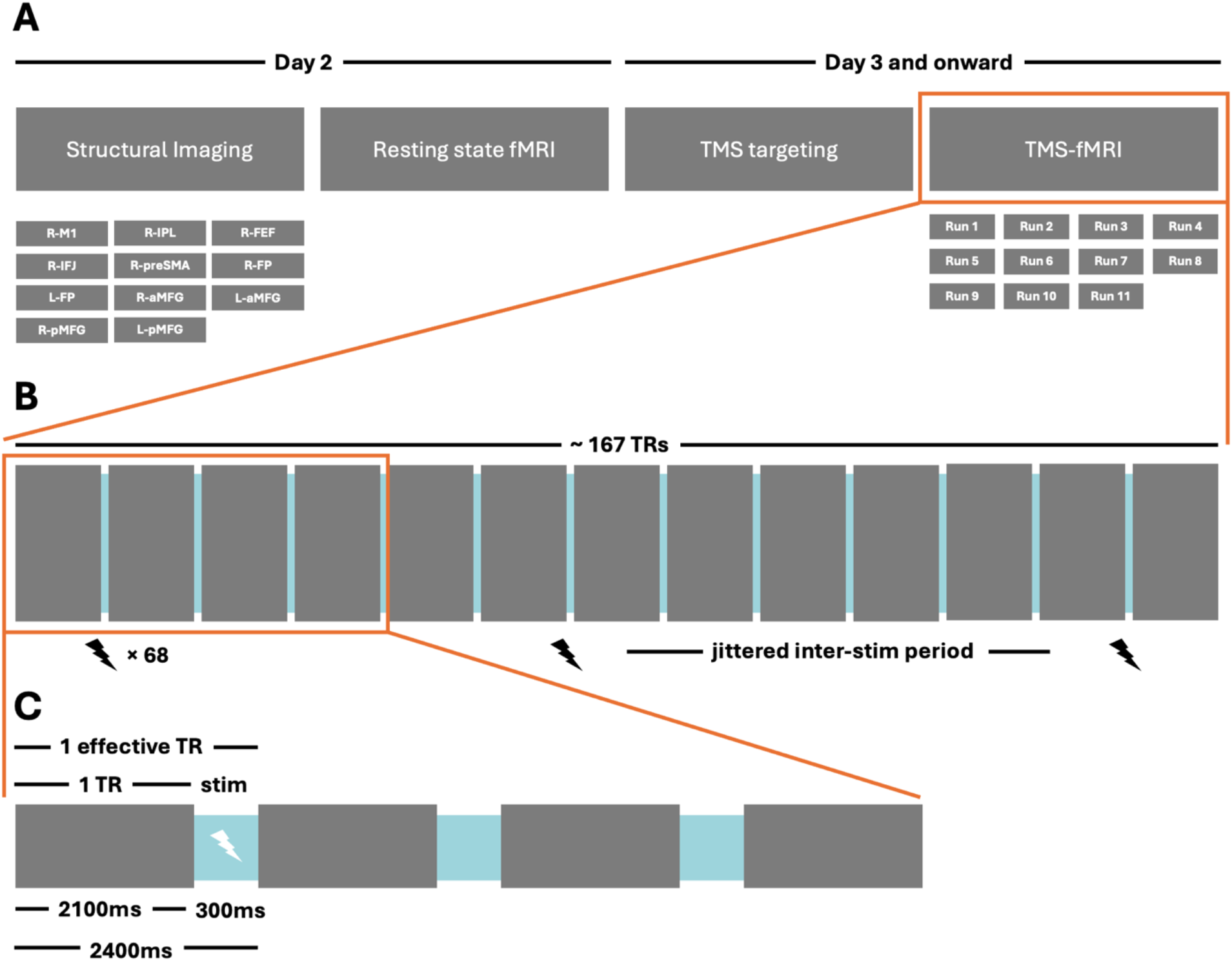
Project timeline and stimulation paradigm at different resolutions. (A) The study consists of 2 sessions, the first being MRI acquisition without TMS and the second including TMS. There were 11 TMS-fMRI runs, each corresponding to an identified cortical region from day 1. (B) The concurrent TMS-fMRI period lasts for ∼167 effective TRs where an effective TR is 2400 ms. (C) 1 Effective TR = 2400 ms where 1 Actual TR = 2100 ms. A 300 ms gap is left in which stimulation may or may not occur. Stimulation occurs on a jittered 6-7 TR basis.

### Anatomical Imaging

All structural scanning was performed at the Stanford School of Medicine Richard M. Lucas Center for Imaging. Images were captured on 3 Tesla GE MR750 scanners with 8-channel head coils. T1 weighted (3D inversion SPGR) images were collected to serve as TMS targeting data for the following sessions. Optimized scan parameters were as follows: acquisition resolution = 1.0 × 0.9 × 0.9 mm, Flip Angle = 15deg, FOV = 22cm, Inversion Time = 450 ms, matrix = 256 × 256, Slices = 184, TE = 3.4ms, TR = 8.6 ms.

### fMRI Data Collection

We report an 8-minute resting state functional Magnetic Resonance Imaging (rs-fMRI) run performed in the same GE MR750 setup. T2-weighted gradient echo spiral in/out pulse sequence (SPRLIO) whole-brain scans with 29 4mm axial slices, Excitations = 2, Flip Angle = 80deg, FOV = 22cm, matrix = 64 × 64, Pixel Size = 3.4mm, Slice Spacing = 0.8mm, TE = 30ms, TR = 2000ms, Volumes = 240 were collected from every participant.

### TMS sites-targeting

We report 11 TMS/fMRI runs, one for each cortical target. Each run consisted of 68 jittered single-pulse TMS doses targeted to a cortical site from the following list: left/right anterior middle frontal gyrus (L/R-aMFG), left/right frontal pole (L/R-FP), left/right posterior middle frontal gyrus (L/R-pMFG), right frontal eye field (R-FEF), right inferior frontal joint (R-IFJ), right inferior parietal lobe (R-IPL), right primary motor area (R-M1), and right pre-supplementary motor area (R-preSMA) (**Table 1, Figure 3B**). The choice of stimulation site locations (MNI coordinates) depended on previous literature (Toll et al. 2020).

**Table 1:**
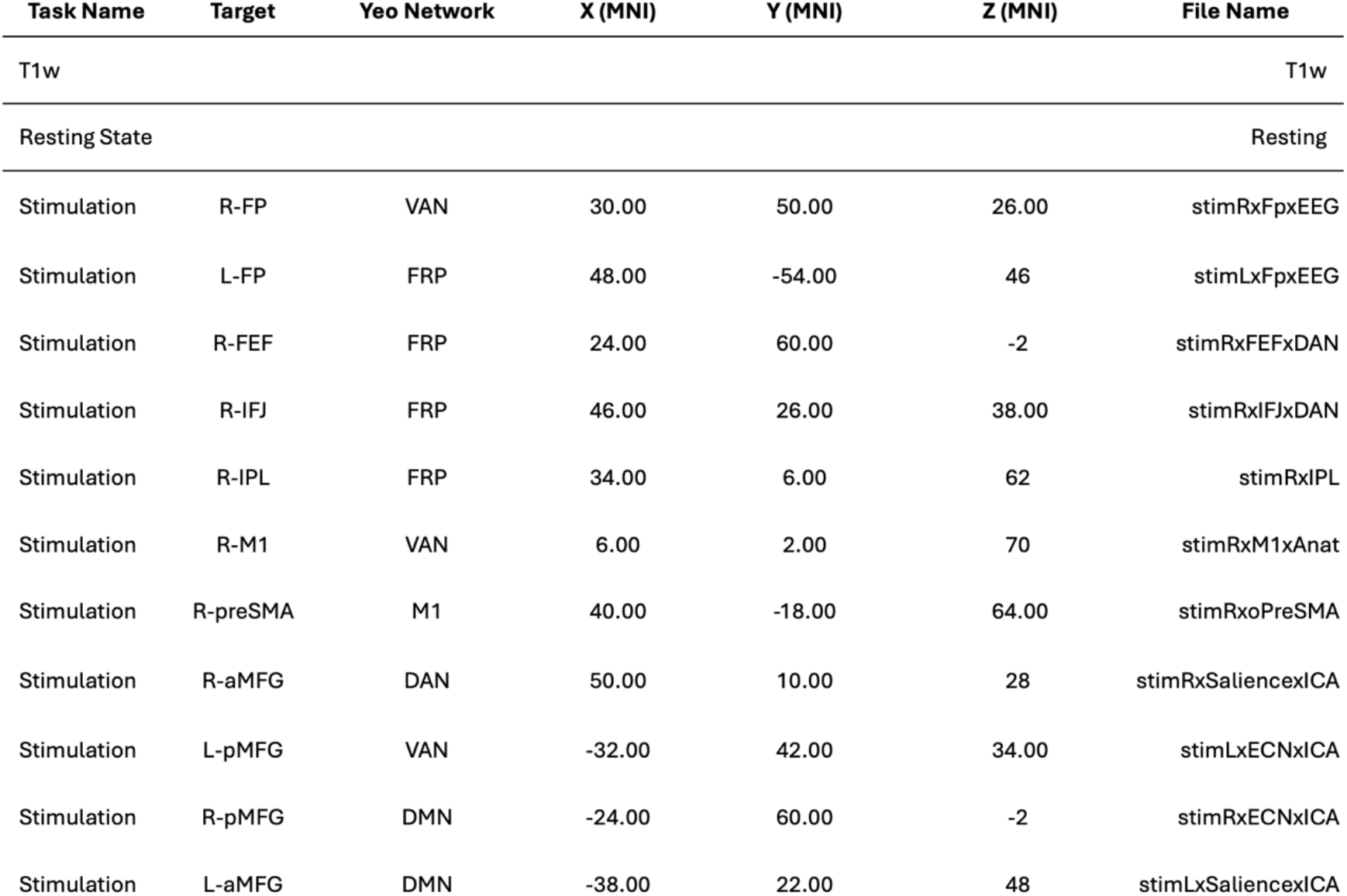
Summary of reported imaging data corresponding to reported stimulation sites.

**Fig 3:**
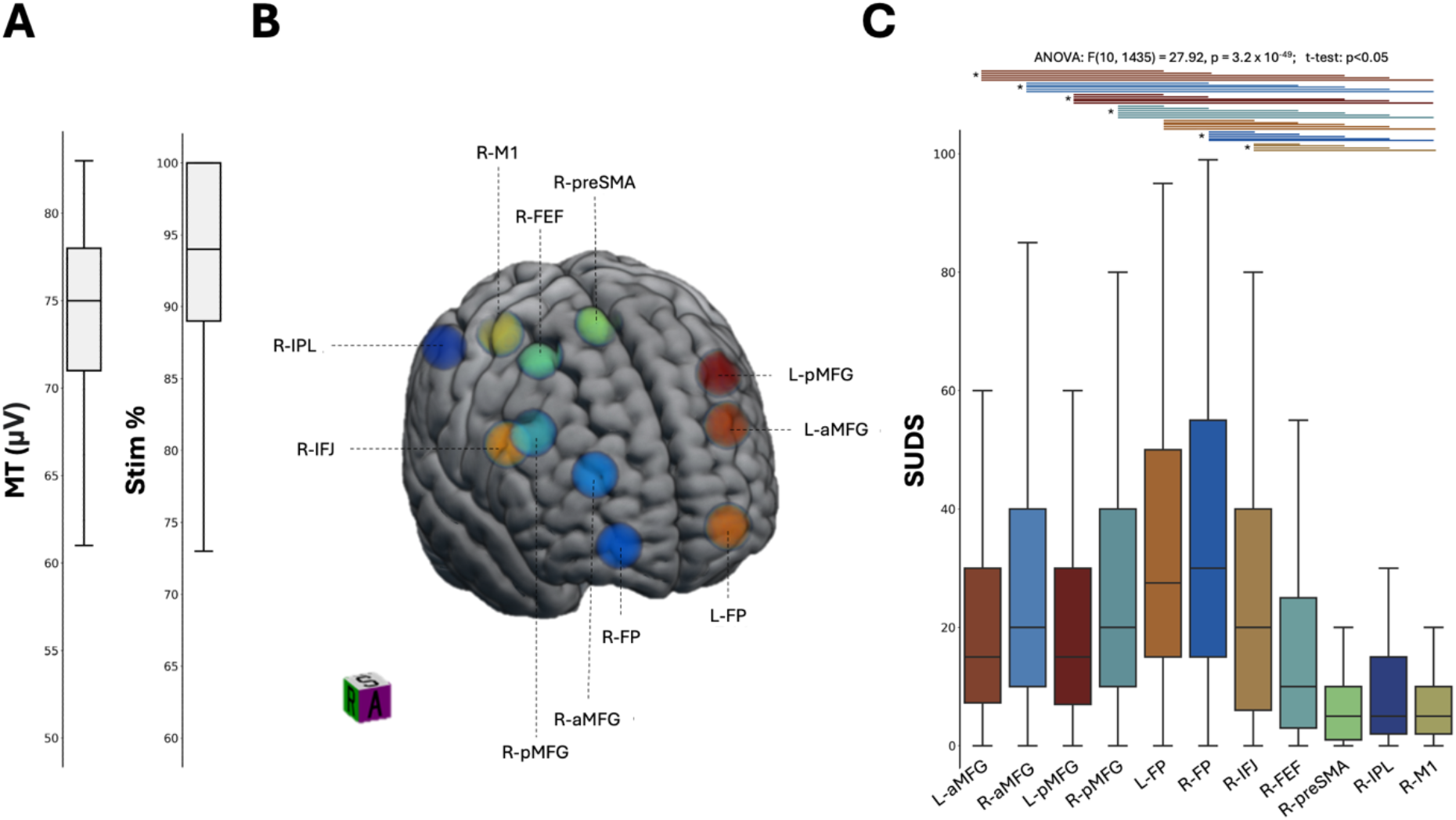
Summary of the stimulation data. (A) Motor Threshold and Stimulation Percentage across all participants. (B) Stimulation sites. (C) Subjective Units of Distress Scale across all participants grouped by stimulation site. Horizontal bars highlight statistical significance (p<0.05) and are colored by stimulation sight.

Prior to TMS/fMRI acquisition, T1 weighted images were used to identify the loci of TMS targets for each participant in the MNI space. These were then co-registered with the head position using the Visor2 software and projected to the individual’s native space. The loci of targeted cortical stimulation were annotated on a Lycra swim cap worn by the individual in the scanner.

### Concurrent TMS/fMRI data collection

A figure-eight, MRI-compatible, TMS coil (MagPro MRi-B91) was set up to deliver biphasic magnetic stimulations. This apparatus was fixed inside the scanner bore via a custom mount and connected to a MagPro X100 stimulator via a patch panel with a powerline filter. Field inhomogeneity resulting from the TMS apparatus was addressed by implementing an automated higher-order shimming optimized for SPRLIO prior to each run. Concurrent TMS/fMRI necessitated using a TMS-compatible single-channel head coil optimized to minimize spatial concerns. MRI parameters mirror those listed in rs-fMRI except for a few key differences. A modified Flip Angle = 85deg, Spacing = 1mm, Volumes = 167 were used. Additionally, an extended TR = 2400ms generated a temporal gap in which single TMS pulses can be interleaved. The TR was set to be 2100ms with a 300ms break, creating an effective TR = 2400ms. Leveraging this break, TMS could be applied without affecting MRI acquisition (**Figure 2B-C**).

Prior to the participant entering the scanner, the Motor Threshold was established using the same TMS apparatus used for concurrent TMS/fMRI collection. Motor threshold and location of maximal stimulation were determined by applying single pulses to the right motor cortex and observing hand twitch responses. MT was defined as the level at which 5/10 consecutive trials resulted in a noticeable motor response in the participant’s hand after pulses to the location of maximal stimulation. All TMS pulses during the TMS/fMRI paradigm were administered at 120% of the established motor threshold (**Figure 3A**).

Each TMS/fMRI run consisted of single pulses with jittered interstimulus intervals ranging from 1 to 6 TR. In total, 68 stimulations were delivered over 6 min 41 secs (max of 167 effective TRs). The TMS apparatus and coil were reset and repositioned for every stimulation site. This procedure was conducted while the participant was still on the table but while the table was extended away from the bore. The order by which each stimulation site was set up and then assessed was randomized to address any confounding order that may be present in the data. Participants were provided with ear protection to minimize the impact of the combined TMS and MRI noise. Additionally, participants wore a GE pulse oximeter and a mid-thoracic strain gauge sampled at 50Hz to measure heart rate and respiration, respectively. Participants were prompted to provide their level of discomfort following every run by asking: “Please rate your pain or discomfort from 1 to 100, ‘0’ meaning ‘no pain’ and ‘100’ meaning ‘the worst pain you can imagine.” (The Subjective Units of Distress (SUDS) scale) (**Figure 3B-C**) Furthermore, participants were notified they could discontinue the experiment at any time. Stimulation to the R-FP was reported as being the most uncomfortable site. ANOVA: F(10, 1435) = 27.92, p = 3.2 × 10^−49^ and t-tests: p<0.05 demonstrate the differences in mean reported SUDS between the stimulation sites (Supplemental Table 1; Figure 3C).

### Data Technical Validation: Assessing Image Quality

All collected data were subdivided using two criteria: Group, e.g., NTHC or TIS, and whether the data was collected during resting state or stimulation. For the stimulation data, the TR was changed to 2400ms post-acquisition to incorporate the 300ms break between the TMS pulses. This was done using the *fslmerge* function from FSL (Woolrich et al., 2009; Smith et al., 2004; Jenkinson et al., 2012). Data was then partially preprocessed via the *mriqc* software package, which provided a host of Image Quality Metrics (IQMs) to describe the data. (Esteban et al. 2017) A subset of IQMs was then selected and reported for data grouped by fMRI paradigm and data grouped by group.

To assess the quality of the MR scans across clinically grouped individuals, participants were broken into 4 categories: NTHC, NTS, TEHC, and TIS. The tSNR, SNR, and mean FD were computed for each group and compared between stimulation and resting state. This demonstrates the similarities and differences between groups and fMRI data acquisition paradigms. Post-hoc t-tests demonstrate that SNR and Mean FD vary between clinical groupings during periods of stimulation (p<0.01) (Figure 4B).

**Fig 4:**
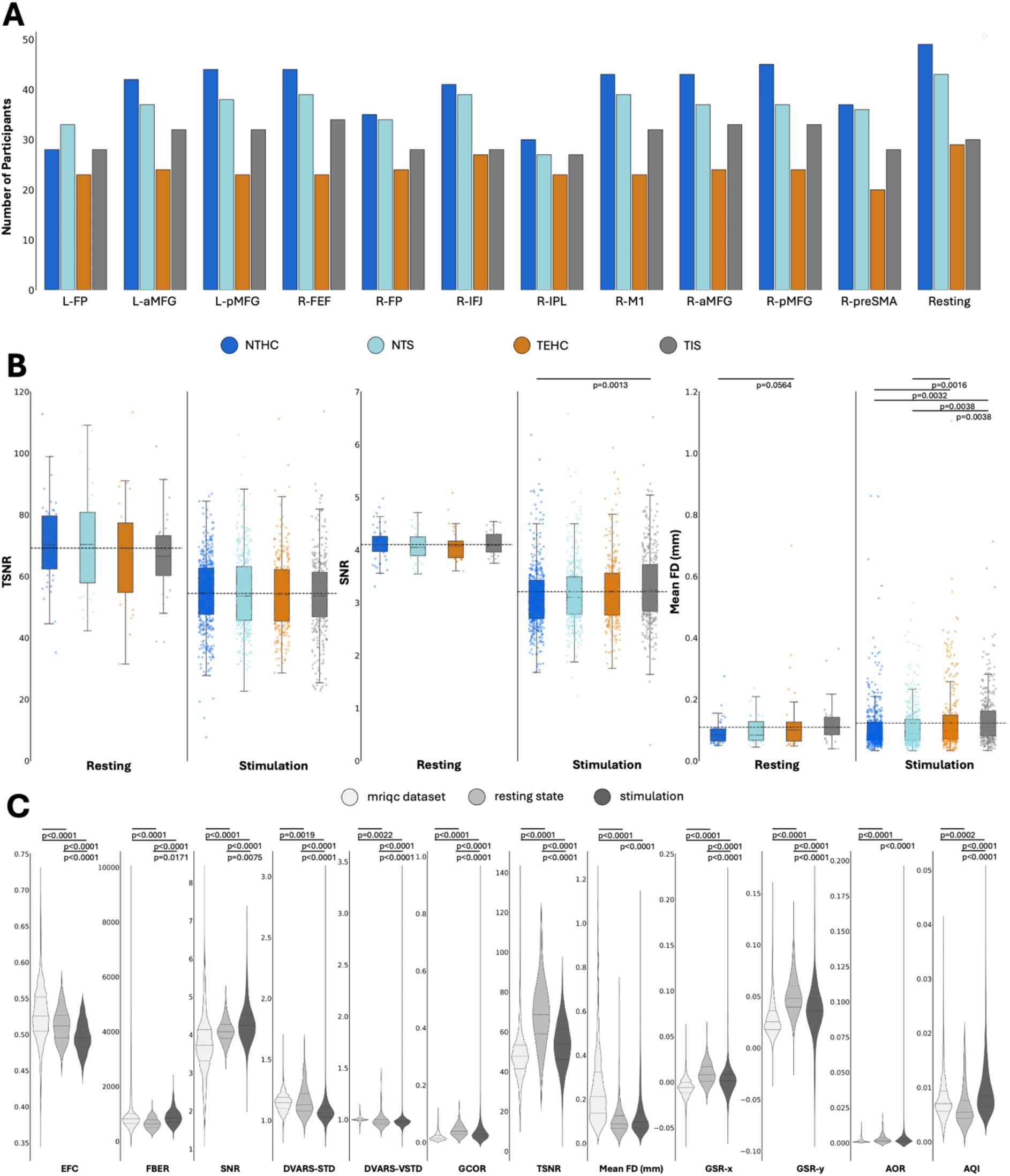
Image Quality Metrics. (A) Number of participants with data grouped by imaging/stimulation target. (B) A comparison of TSNR, SNR, and Mean FD across clinical groupings. The Dashed line represents the mean across groups. Horizontal bars highlight statistical significance. (C) A comparison of group level Image Quality Metrics as compared to the mriqc database deplete of outliers greater than 5 IRQs from mean. Horizontal bars highlight statistical significance.

For data grouped by fMRI paradigm (resting or TMS), we report Spatial Metrics: Entropy-focus criterion (EFC; Atkinson et al. 1997), Foreground-Background energy ratio (FBER; Shehzad et al. 2015), Signal-to-Noise Ratio (SNR); Temporal Metrics: DVARS (D referring to the temporal derivative of time courses, VARS referring to RMS variance over voxels; Power et al. 2012), Global Correlation (GCOR; Saad et al. 2013), TSNR (Temporal SNR; Kruger et al. 2001); and Artifact Metrics: Framewise Displacement (FD; Jenkinson et al. 2002), Ghost to Signal Ratio (GSR; Giannelli et al. 2010), AFNI’s outlier ratio (AOR), and AFNI’s quality index (AQI). These IQMs were compared to the open source *mriqc* dataset. (Esteban et al. 2019) The *mriqc* database was scrubbed of outliers 5 times the interquartile distance away from the mean for easy plotting and comparison to the collected dataset. The collected data was not modified. While the data quality between the aforementioned data and established fMRI data remain quite similar, p-values derived from post-hoc t-tests demonstrate the differences in each measure. EFC, FBER, SNR, DVARS, TSNR, and Mean FD from the collected data outperform metrics from the mriqc dataset (Figure 4C).

### Dissemination of Data

The aforementioned de-identified data were posted and will be periodically updated on the OpenNeuro website (Markiewicz et al. 2021). The data are organized via the Brain Imaging Data Structure (BIDS) version (v1.9.0 guidelines (Gorgolewski, K., Auer, T., Calhoun, V. et al., 2016). As the BIDS guidelines dictate, data has been organized into nested files with standardized names. Acquired data are converted to NIFTI format with associated metadata stored in JSON files. Subjects are organized into four categories: NTHC, NTS, TEHC, and TIS (**Figure 1A-B**).

## Data Records

The dataset comprises 152 subjects categorized into four groups: NTHC, TEHC, NTS, and TES, with IDs in the 1000s, 2000s, 3000s, and 4000s, respectively (**Figure 1C**). All data has been de-identified. The labeling of the data is as follows:

Structural data: sub-XXXX_ses-Y_run-Z_T1w.ext,

Functional data: sub-XXXX_ses-Y_task-TYPE.ext,

where XXXX is the subject ID, Y is the session number, Z indicates the run number, TYPE refers to the type of fMRI scan (either resting or stimulation), and *ext* represents the file extension (.nii, .nii.gz, or .json).

For stimulation tasks, the naming convention also specifies the hemisphere, the specific stimulation site out of the 11 available, and the associated brain network. For example, “stimRxFEFxDAN” denotes stimulation of the right hemisphere’s Frontal Eye Field (FEF), associated with the Dorsal Attention Network (DAN).

## Technical validation

To assess the quality of the collected data, we took a three-step approach. First, we assessed the quantity. Second, we hierarchically grouped the data to tease out differences between clinical groups and between resting state fMRI and TMS-fMRI. Finally, we compared the collected data with the *mriqc* public dataset.

### Participants and Data Quantity

333 healthy participants (204M, 127F, 2 Unreported) were recruited with a mean age of 32 years (SD: 11). Of these 333 recruited participants, we report data from 152 individuals (93M, 52F, Mean age 31.1yr, SD age 10.5yr) (**Figure 1C)**. The quantity of present data differs between groups (**Figure 1C**). When examining the number of reported participants assigned to individual groups, NTHC reports the largest number of participants at 46, and TIS reports the lowest number of participants at 34. On average, NTHC group consistently reports the most data across both resting-state and TMS/fMRI, with a total completion rate of 78.6% across all individuals. TEHC group reports the lowest total completion at 47.8% across all individuals. Differences in data completion are attributed to participant dropouts and/or incompletion of all 11 TMS targets (**Figure 4A**).

### Functional Data Quality

When compared between groups, the reported IQMs (TSNR, SNR, Mean FD) remain consistent (**Figure 4B**). This demonstrates the robustness of the scan protocol to varying healthy and clinical populations. Furthermore, the lack of variance in average IQMs across clinical groups allows future valid comparisons between sub-groups. Resting-state fMRI average TSNR and SNR outperformed TMS/fMRI IQMs at the clinical subgroup and group levels. This is to be expected as introducing TMS apparatus into the scanner bore impacts scan quality. Notably, the mean FD during stimulation was slightly higher than the mean FD during rest. This could be due to the added stress of the TMS apparatus or the added pressure on participants to remain still during stimulation. Despite the reduction in scan quality during TMS/fMRI, all reported IQMs for all clinical groups and scanning paradigms stay within the acceptable ranges.

### Functional Data Compared to Canonical Data

To assess the quality of our collected fMRI data, we first split the data into two subgroups: rfMRI and TMS/fMRI. These two groups were then compared to the *mriqc* published dataset via key IQMs (**Figure 4C**). Our collected data remained similar to the published dataset, so we believe the data maintains a quality consistent with previously published fMRI datasets.

## Limitations and General Usage Notes

The data were organized using the BIDS naming convention, employing key-value pairs such as “sub-value,” “run-value,” and “task-value.” The “sub” key represents study participants. The “run” key denotes separate T1-weighted (T1w) data acquisitions for the same subject during the same session using consistent parameters. The “task” key is used for BOLD (Blood Oxygenated Level Dependent) fMRI data and identifies either resting-state activity or one of the 11 stimulation sites detailed in the methods section.

Most participants had their structural and resting-state data collected in one session, with stimulation data gathered in a separate session. However, some participants underwent more than two sessions for data collection. In cases where structural and resting-state data were not collected during the first session, these participants may have their data recorded in subsequent sessions, and session one may be absent for them.

To ensure anonymization, the structural data was defaced using PyDeface (https://pypi.org/project/pydeface/), as recommended by OpenNeuro.

## Code/data availability

All data associated with the study are posted at Open Neuro under the name (OpenNeuro Dataset ds005498, https://openneuro.org/datasets/ds005498). All processing and QC code is posted to the Brain Dynamics Lab GitHub https://github.com/braindynamicslab/sptmsfmri

## Acknowledgements

This project was funded by an NIH R01 MH103324 awarded to A.E.

## Figures and Tables

**Supplemental Table 1:** Summary of significant pairwise comparisons between reported SUDS grouped by region.

| Region 1 | Region 2 | P-value | Region 1 | Region 2 | P-value | Region 1 | Region 2 | P-value |
| --- | --- | --- | --- | --- | --- | --- | --- | --- |
| L-aMFG | L-FP | 0.0001 | R-pMFG | L-FP | 0.0144 | R-FP | R-IFJ | 0.0461 |
| L-aMFG | R-FP | <0.0001 | R-pMFG | R-FP | 0.0002 | R-FP | R-FEF | <0.0000 |
| L-aMFG | R-preSMA | <0.0001 | R-pMFG | R-FEF | 0.0049 | R-FP | R-preSMA | <0.0000 |
| L-aMFG | R-IPL | 0.0002 | R-pMFG | R-preSMA | <0.0000 | R-FP | R-IPL | <0.0000 |
| L-aMFG | R-M1 | <0.0001 | R-pMFG | R-IPL | <0.0000 | R-FP | R-M1 | <0.0000 |
| R-aMFG | R-FP | 0.0030 | R-pMFG | R-M1 | <0.0000 |  |  |  |
| R-aMFG | R-FEF | 0.0003 | L-FP | R-IFJ | 0.0022 |  |  |  |
| R-aMFG | R-preSMA | <0.0001 | L-FP | R-FEF | <0.0001 |  |  |  |
| R-aMFG | R-IPL | <0.0001 | L-FP | R-preSMA | <0.0001 |  |  |  |
| R-aMFG | R-M1 | <0.0001 | L-FP | R-IPL | <0.0001 |  |  |  |
| L-pMFG | L-FP | 0.0001 | L-FP | R-M1 | <0.0001 |  |  |  |
| L-pMFG | R-FP | <0.0001 | R-IFJ | R-FEF | 0.0461 |  |  |  |
| L-pMFG | R-preSMA | <0.0001 | R-IFJ | R-preSMA | <0.0001 |  |  |  |
| L-pMFG | R-IPL | 0.0003 | R-IFJ | R-IPL | <0.0001 |  |  |  |
| L-pMFG | R-M1 | 0.0001 | R-IFJ | R-M1 | <0.0001 |  |  |  |

## References

Jenkinson, M., Beckmann, C. F., Behrens, T. E. J., Woolrich, M. W. & Smith, S. M. FSL. Neuroimage 62, 782–790 (2012).

Smith, S. M. et al. Advances in functional and structural MR image analysis and implementation as FSL. Neuroimage 23 Suppl 1, S208–19 (2004).

Woolrich, M. W. et al. Bayesian analysis of neuroimaging data in FSL. Neuroimage 45, S173–86 (2009).

Power, J. D. et al. Functional network organization of the human brain. Neuron 72, 665 (2011)

Deco, G., Jirsa, V. K. & McIntosh, A. R. Emerging concepts for the dynamical organization of resting-state activity in the brain. Nature Reviews Neuroscience 2011 12:1 12, 43–56 (2010).

Cole, M. W., Ito, T., Bassett, D. S. & Schultz, D. H. Activity flow over resting-state networks shapes cognitive task activations. Nat. Neurosci. 19, 1718–1726 (2016).

Miŝic, B. et al. Network-Level Structure-Function Relationships in Human Neocortex. Cereb. Cortex 26, 3285–3296 (2016).

Gollo, L. L., Roberts, J. A. & Cocchi, L. Mapping how local perturbations influence systems-level brain dynamics. Neuroimage 160, 97–112 (2017).

Bestmann, S. et al. Mapping causal interregional influences with concurrent TMS–fMRI. Exp. Brain Res. 191, 383–402 (2008).

Ruff, C. C., Driver, J. & Bestmann, S. Combining TMS and fMRI: From ‘virtual lesions’ to functional-network accounts of cognition. Cortex 45, 1043–1049 (2009).

Esteban, O. et al. Crowdsourced MRI quality metrics and expert quality annotations for training of humans and machines. Scientific Data 2019 6:1 6, 1–7 (2019).

Atkinson, D., Hill, D. L. G., Stoyle, P. N. R., Summers, P. E. & Keevil, S. F. Automatic correction of motion artifacts in magnetic resonance images using an entropy focus criterion. IEEE Trans. Med. Imaging 16, 903–910 (1997).

Power, J. D., Barnes, K. A., Snyder, A. Z., Schlaggar, B. L. & Petersen, S. E. Spurious but systematic correlations in functional connectivity MRI networks arise from subject motion. Neuroimage 59, 2142 (2012).

Saad, Z. S. et al. Correcting brain-wide correlation differences in resting-state FMRI. Brain Connect. 3, 339–352 (2013).

Triantafyllou, C., Polimeni, J. R. & Wald, L. L. Physiological noise and signal-to-noise ratio in fMRI with multi-channel array coils. Neuroimage 55, 597–606 (2011).

Jenkinson, M., Bannister, P., Brady, M. & Smith, S. Improved Optimization for the Robust and Accurate Linear Registration and Motion Correction of Brain Images. Neuroimage 17, 825–841 (2002).

Giannelli, M., Diciotti, S., Tessa, C. & Mascalchi, M. Characterization of Nyquist ghost in EPI-fMRI acquisition sequences implemented on two clinical 1.5 T MR scanner systems: effect of readout bandwidth and echo spacing. J. Appl. Clin. Med. Phys. 11, 170–180 (2010).

Markiewicz, C. J. et al. The openneuro resource for sharing of neuroscience data. Elife 10, (2021).

Gorgolewski, K. J. et al. The brain imaging data structure, a format for organizing and describing outputs of neuroimaging experiments. Scientific Data 2016 3:1 3, 1–9 (2016).

Toll, R. T. et al. An electroencephalography connectomic profile of posttraumatic stress disorder. Am. J. Psychiatry 177, 233–243 (2020).

